# A Method to Calibrate Chemical-agnostic Quantitative Adverse Outcome Pathways on Multiple Chemical Data

**DOI:** 10.1101/2025.03.11.642550

**Authors:** Zheng Zhou, Ullrika Sahlin

## Abstract

Quantitative Adverse Outcome Pathways (qAOPs) may support next-generation risk assessment by integrating New Approach Methodologies (NAMs) for derivation of points of departure. To be useful, a qAOP should be chemical-agnostic. However, existing qAOP studies often pool multi-chemical data without adequately addressing inter-chemical heterogeneity. Consequently, fundamental pathway relationships become obscured by heterogeneity-induced noise, thereby compromising the reliability of chemical-agnostic predictions. We developed a chemical-agnostic calibration approach to addresses this challenge by leveraging hierarchical structures to systematically separate chemical-specific heterogeneity from underlying pathway effects. Through this methodological framework, chemical-specific deviations are explicitly modeled as random effects, enabling the extraction of pathway-level parameters that represent core mechanistic relationships independent of individual chemical properties. Through simulation studies across varying heterogeneity levels, we demonstrate that performance differences between models with and without hierarchical calibration reveal the magnitude of heterogeneity in the data. Moreover, when heterogeneity is substantial, an uncalibrated qAOP should not be considered truly chemical-agnostic in practice, as it confounds pathway-level effects with chemical-specific variation. We demonstrated the application of this calibration approach through a case study of non-mutagenic liver tumor. The framework proposed in this study enhances qAOP generalizability while preserving the chemical-agnostic principle, supporting robust NAMs-based next-generation risk assessments.

Figure 1: For TOC only

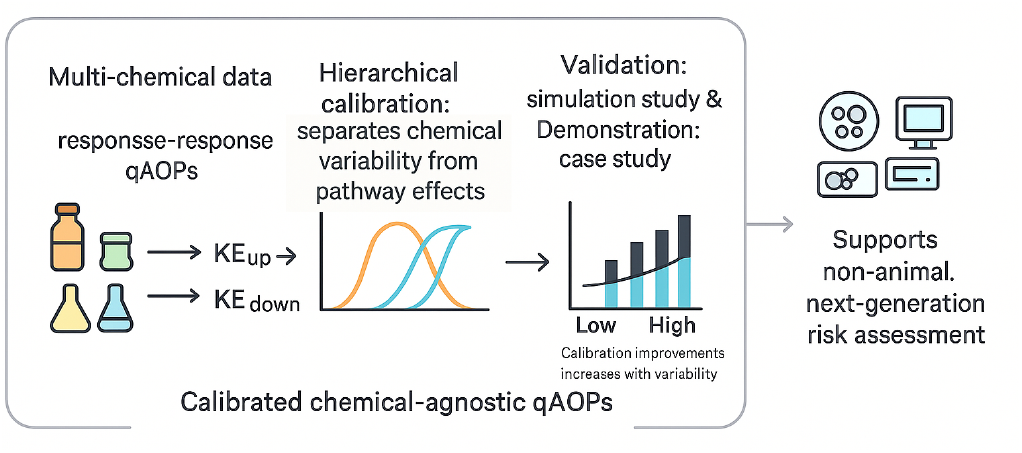

## 1 Introduction

### 1.1 Background

The Adverse Outcome Pathway (AOP) framework organizes existing knowledge on biological mechanisms that link molecular or cellular perturbations to adverse outcomes (AOs) across different levels of biological organization. ^1^ It facilitates the extrapolation of mechanistic evidence from animal and non-animal models at lower biological levels to predict apical AOs. ^2,3^ Quantitative AOPs (qAOPs) represent an advanced stage of AOP development and application, focusing on the quantitative relationships among key events (KEs) and between KEs and AOs. ^2–4^ KEs are measurable biological changes that are necessary, but not sufficient, for the occurrence of an AO. ^1^ Robust AOPs and qAOPs are essential for implementing New Approach Methodologies (NAMs) aimed at the 3Rs—Reduction, Replacement, and Refinement—of animal testing in next-generation risk assessments. ^5^ qAOPs have been increasingly applied to support hazard identification, derive points of departure (PODs), and develop integrated testing and assessment approaches for regulatory decision-making. ^4,6^

### 1.2 Challenge to Achieve Chemical Agnostic Adverse Outcome Pathways

Chemical agnosticism represents a fundamental principle of AOPs and qAOPs, ^1^ effectively distinguishing them from other mechanistic frameworks such as mechanisms of action (MOAs). ^7,8^ This distinction manifests in two key aspects. First, AOPs initiate at the molecular initiating event, whereas MOAs are dose-dependent and begin with chemical dose or exposure. Second, AOPs operate independently of specific chemicals; provided that the initial event is triggered with sufficient magnitude, the downstream chain of key events can be activated regardless of the causative agent. By contrast, MOAs remain intrinsically linked to specific chemical exposures and their dose-response relationships.

However, existing qAOP studies exhibit limited adherence to the chemical-agnostic principle in two important ways. First, several studies have constructed preliminary models using data from a single chemical and a specific exposure route or measurement assay. ^9,10^ Although these models may demonstrate excellent performance for the specific chemical examined, their validity for the inference of other chemicals leading to the same AO remains unverified. This lack of generalizability significantly restricts the applicability of such qAOP models for prospective prediction of new chemicals in regulatory and industrial contexts. Second, some qAOP studies have developed models using multi-chemical datasets while inadequately addressing cross-chemical heterogeneity. ^11–13^ Such heterogeneity arises from various factors, including differences in chemical structures and properties, ^1^ which consequently lead to substantial variation in the potency required to trigger upstream key events across chemicals. When heterogeneity is pronounced, a dose sufficient to induce the complete cascade of key events and the adverse outcome in one chemical may fail to trigger the initial event in another. Therefore, neglecting this heterogeneity undermines the fundamental chemical-agnostic principle and compromises model reliability across diverse chemical exposures. When data exhibiting substantial cross-chemical heterogeneity are pooled without accounting for such differences, the pooled model would differ significantly from the models derived on individual chemicals and may be biased by extreme data points. ^14^

### 1.3 Study Objectives

Given the importance to consider heterogeneity in multi-chemical datasets, it is essential to distinguish chemical-specific heterogeneity from the underlying chemical-agnostic pathway when developing and applying qAOPs. Hierarchical modeling is an established statistical parametric approach for considering grouped data ^14,15^ which has been successfully applied to dose-response modeling. ^16,17^ The implementation of hierarchical structures in qAOP models, particularly parametric response-response models, remains largely unexplored. By incorporating hierarchical frameworks into qAOP development, it is possible to explicitly account for cross-chemical variability while preserving the chemical-agnostic principles fundamental to the AOP framework. This study addresses this methodological gap by developing and evaluating an hierarchical response-response model for multi-chemical qAOP applications.

The aim of this study is to develop a framework to calibrate chemical-agnostic qAOP models to facilitate more robust and generalizable qAOPs. Specifically, the objectives are to: 1) promote the incorporation of multi-chemical data in qAOP development; 2) develop a modeling approach that can effectively separate chemical-specific heterogeneity from chemical-agnostic pathways using hierarchical structures; 3) evaluate the predictive performance of the qAOP with hierarchical structures against the one without.

## 2 Theory and methods

### 2.1 Quantitative Adverse Outcome Pathway Model

The following qAOP model is considered for the chemical-agnostic calibration of this study: Observed values of *KE*_*up*_ are continuous, denoted as a random variable *Y*_1_; Observed values of *KE*_*down*_ are dichotomous, denoted as a random variable *Y*_2_. The Benchmark Dose (BMD) methodology is employed to model the dose-response relationship between dose and *Y*_1_ and the response-response relationship between *Y*_1_ and *Y*_2_. The assumption from BMDS ^18,19^ are followed as in Equation (1): 1) The outcome of *KE*_*up*_ follows a lognormal distribution at any doses *d*, where the expected value of *Y*_1_ on the log scale is dependent on dose *d* and parameters *θ*_1_ according to a function *f*_1_(*d*|*θ*_1_); 2) The outcome of *KE*_*down*_ follows a Bernoulli distribution at any level of *Y*_1_, where the probability of *Y*_2_ is dependent on *Y*_1_|*d* and parameters *θ*_2_.

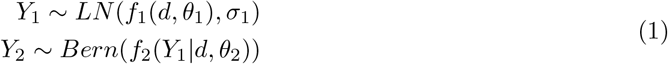

The log likelihood function for the parameters *θ*_1_ is:

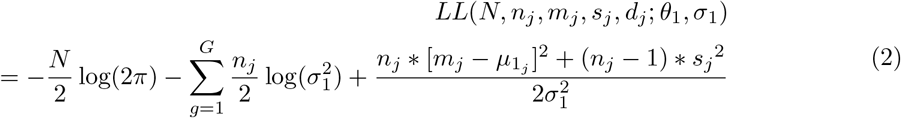

where *N* is total number of observations, *j* = 1, …, *J* is the dose group, *d*_*j*_ is the dose, *n*_*j*_ is the number of subjects, *m*_*j*_ is the sample mean and *s*_*j*_ is the sample standard deviation for group *j*. 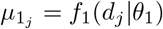 is the expected response for group *j*.

Denoting the probability for the outcome *KE*_*down*_ in group *j* as *p*_*j*_ = *f*_2_(*Y*_1_|*d*_*j*_, *θ*_2_), the log likelihood for the parameters *θ*_2_ is:

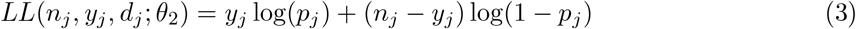

where *j* = 1, …, *J* is the dose group (note that the dose groups can alter between data sets), *n*_*j*_ is the sample size, *y*_*j*_ the number of outcomes, and *d*_*j*_ the dose for group *j*. For details on the application of Benchmark dose modelling, we recommend the User Guide ^20^ and Technical Guidance ^18^ of U.S. EPA Benchmark Dose Software (BMDS) and EFSA guidance. ^19^

For the dose-response to the first key event *f*_1_, we used a continuous-Hill function from BMDS ^18^ (Equation (4)) with parameters *θ*_1_ = {*a, b, c, g* }, where *a* is background response; *b* is maximum change, *b >* 0 & *b <* 0 for increasing and decreasing trends respectively; *c >* 0 is the dose at which half-maximal change occurs; *g >* 0 is power, i.e., the steepness of sigmoid curves. A half-maximal potency parameter is the level of the dose leading to half of the observed maximum change in the response variable.

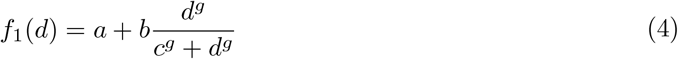

For the response-response part, a dichotomous-Hill function from BMDS ^18^ is adapted for *f*_2_ (Equation (5)) with parameters *θ*_2_ = *v, q, h, r*. where 0 *< v* ≤ 1 is maximum probability; 0 ≤ *q <* 1 is background risk so that *vq* is background probability; *h* is half-maximal potency; *r* ≥ 0 is power, i.e., steepness of sigmoid curves.

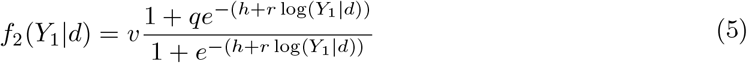

The Hill functions are used because they are flexible and biologically relevant. ^18,21^ The flexibility of Hill functions allows the modeling of a potential threshold in the dose-response and response-response relationship.

### 2.2 Hierarchical Structure

For each chemical *k* = 1, …, *K* there are *N*_*k*_ observations. Data may be reported in summarized format that the number of subjects and summary statistics per group are available, such that data are indexed with *i* = 1 … *J*_*k*_, where *J*_*k*_ is the number of dose groups per chemical.

A “*flat*” response-response model, i.e. without hierarchical structure is the dichotomous-Hill function in Equation (5) with the same set of parameters *θ*_1_ = {*v, q, h, p*} for all chemicals. The hierarchical structure incorporates two key features (Equation (6)). First, parameters are assigned to hierarchical levels based on their mechanistic interpretation: parameters *v* and *q* are chemical-specific, while *h* and *r* are chemical-agnostic. This designation of *h* and *r* as chemical-agnostic is justified on two grounds. From a mechanistic perspective, these parameters control the location and steepness of the sigmoid region of the response-response curve, respectively, thereby reflecting the critical threshold levels of *KE*_*up*_ required to induce *KE*_*down*_. These threshold characteristics are inherent to the biological pathway rather than specific chemicals. Moreover, empirical evidence from previous meta-dose-response studies demonstrates that estimated log-inflection points and steepness parameters exhibit minimal variation across chemicals, even when modeled individually. ^22,23^ Consequently, maintaining (h) and (p) as chemical-agnostic parameters is both mechanistically justified and mathematically efficient. Second, heterogeneity is parameterized through a combination of random and fixed effects. Specifically, the structure of each hierarchical parameter comprises three components (Equation (6)): (a) a chemical-agnostic central location, (b) chemical-agnostic variability, and (c) chemical-specific relative deviations from the central location. The product of the chemical-agnostic variability and chemical-specific relative deviations constitutes the random effects that capture departures from the central pathway for each chemical.

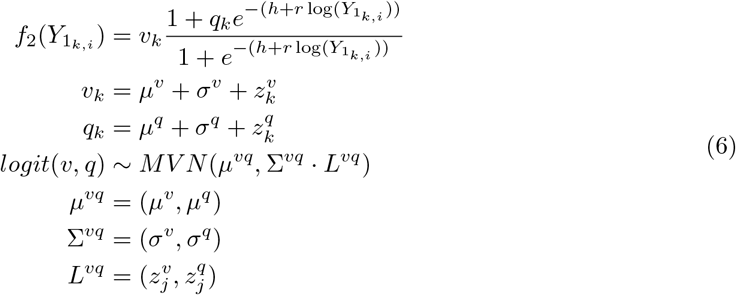

where a superscript indicates the dependent, low-level parameter; *µ* and *σ* are chemical-agnostic central location and variability respectively; *z*_*k*_ are chemical-specific relative deviations; MVN refers to multivariate normal distributions, which depends on a mean vector *µ*, a standard deviation matrix Σ and a correlation matrix *L*.

Identical parameterization is employed for the dose-response component of both flat and hierarchical models to control for uncertainty arising from dose-response modeling. Notably, this parameterization is not necessarily chemical-specific, ensuring that both model structures share the same dose-response assumptions. By maintaining consistency in the dose-response modeling approach, any differences in predictive performance between the flat and hierarchical models can be attributed solely to structural differences in the response-response calibration. This design allows for direct comparison of model performance and isolates the impact of incorporating hierarchical structures. In summary, the flat qAOP includes a chemical-specific dose-response part and a flat response-response part, while the hierarchical qAOP has a chemical-specific dose-response part and a hierarchical response-response part.

### 2.3 Model performance

The impact of the hierarchical calibration approach is evaluated by comparing the predictive performance of flat to hierarchical model specification. Model performance is assessed using the expected log predictive density (ELPD, Equation (7)), which enables comparison of posterior predictive performance between non-nested, non-linear models, ^24,25^ and the Widely Applicable Information Criterion (WAIC), ^25^ to check the robustness towards the choice of metric. The ELPD is defined as

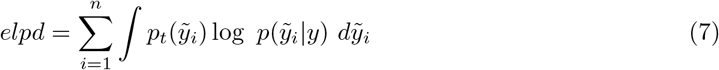

where 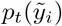 is the distribution for the true data-generating process for 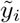. In this study, 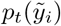 is approximated by Pareto Smoothed Importance Sampling Leave-one-out Cross Validation (PSIS-LOO-CV), ^26^ implemented using the ‘*loo*’ package in R. ^27^ For brevity, *elpd*_*P SIS* ^−^ *LOO*_ is hereafter referred to as LOO. According to the definition of LOO, ^25,26^ if *LOO*_*hierarchical*_ exceeds *LOO*_*flat*_, the hierarchical model demonstrates superior predictive performance compared to the flat model. Such a result indicates that cross-chemical heterogeneity in the data is sufficiently pronounced to warrant the use of hierarchical structures. Conversely, if the two models exhibit comparable LOO values, the added complexity of the hierarchical structure may not be justified for the dataset under consideration.

### 2.4 Data Requirements

To characterise the effect of dose on both upstream and downstream key event responses the chemical-agnostic calibration framework requires (1) two dose-response datasets corresponding to the upstream and downstream key events, respectively, and (2) some chemicals shared between these two datasets. High-quality data are essential for robust model development and validation. The highest quality toxicological data originate from randomized controlled trials, which provide unbiased estimates of dose-response relationships. ^6^ When experimental data are unavailable, data from observational studies may be considered as an alternative. However, observational data carry substantial risks of bias, resulting in lower weight of evidence ^6,7^ and potentially compromising the robustness of the qAOP framework. To enable meaningful characterization of cross-chemical heterogeneity, both datasets must include multiple shared chemicals (and not just only one).

Data quality can be further enhanced when both datasets originate from the same experiment, they naturally share the same applicability domain (such as an *in vivo* route of exposure or an *in vitro* assay) and dosage specifications (i.e., dose groups), thereby ensuring optimal comparability. Conversely, when the two datasets are collected from separate experiments or studies, the shared chemicals must have identical or at least comparable applicability domains to maintain validity. In such cases, complete alignment of dosage setups across the original datasets is not strictly necessary as long as doses are converted to consistent units compatible within the qAOP framework. Calibration of dose-response models can be done with individual-level response data as well as sufficient summary statistics thereof. For performance evaluation, it is best to have access to individual-level data, since aggregation results in information loss. Preserving the highest resolution of data whenever possible enhances the reliability and precision of the hierarchical calibration approach.

### 2.5 Model Implementation

Model calibration was done through Bayesian updating using a No-U-Turn Sampler in Hamiltonian Monte Carlo sampling through the R interface of Stan. ^28^ The target log-density functions and complete Stan scripts are available in the supplementary information. Stan scripts for both flat and hierarchical models are provided in the supplementary information.

Weakly informative priors were assigned to all model parameters to limit prior influence while maintaining computational stability 1. In particular, we avoid using uniform distributions for priors, motivated by that for some chosen ranges on benchmark dose parameter this can result in bias in the derived point of departure. ^29^

There are ways to specify hyper parameters by comparing the prior predictive distribution to what is reasonable over observations. ^30^ Here, probability distributions for priors were chosen by reasoning and hyper parameter within these models specified using a simple iterative prior predictive evaluation procedure to ensure they yielded plausible predictions within the context of tumor incidences. A set of hyperparameter values was initially drawn from the specifications of the source study, and the corresponding prior predictive distribution was generated. The resulting distribution was evaluated by comparing its range and shape against the empirical observations of tumour incidences, which were bounded between 0 and 1. Hyperparameters were iteratively adjusted until the prior predictive distribution adequately spanned the full data range and exhibited realistic variability without excessive concentration in any subregion of the response space. The final set of hyperparameter values used in the analyses is provided in the Supplementary Information.

The MCMC sampling procedure comprises four chains, each consisting of 5,000 warm-up iterations and 5,000 sampling iterations. The performance of the MCMC samplers is evaluated for chain convergence and sampling efficiency using the 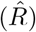 statistic and effective sample size (ESS), respectively. Convergence is deemed satisfactory when 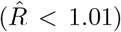 for all parameters, indicating that the chains have adequately explored the posterior distribution. Additionally, adequate sampling efficiency is confirmed when ESS exceeds 400 for all parameters, ensuring sufficient independent samples for reliable posterior inference.

### 2.6 Simulation Study

A simulation study was conducted to validate the calibration framework across varying levels of cross-chemical heterogeneity. The simulation procedure is illustrated in Figure 1. To isolate the impact of hierarchical calibration on the response-response component, both flat and hierarchical models employ identical dose-response models, thereby ensuring that any performance differences can be attributed solely to the response-response calibration structure. Complete scripts for the simulation study and subsequent analyses are provided in the supplementary information.

**Figure 1.**
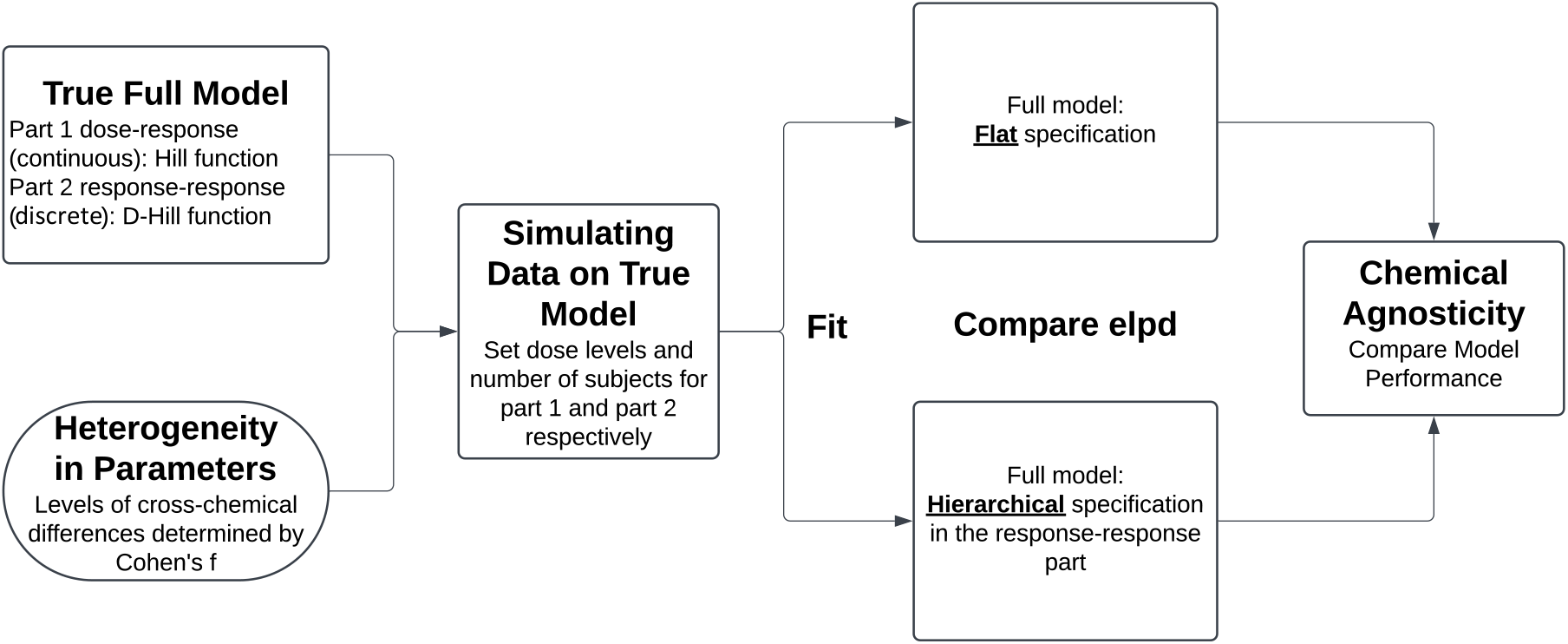
Diagram of Simulation Study

Datasets for the simulation study were generated at four levels of cross-chemical heterogeneity (none, small, medium, and large) by increasing the variance of the random terms in the hierarchical model. To verify heterogensity levels, the ratio of cross- and within-chemical variance, adjusted for doses, was calculated as Cohen’s f effect size ^31^ following Equation (8). A large effect size indicates that the changes in *KE*_*down*_ across chemicals are bigger than that of the same chemical over *KE*_*up*_.

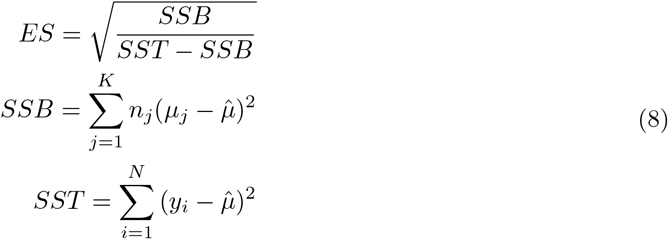

where SSB and SST are between-chemical and total squared variance, respectively; 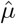 is global mean of the data; j is chemical index, *n*_*j*_ and *µ*_*j*_ are the size and mean per chemical. Response-response parameters were iteratively adjusted until the computed Cohen’s f values of the simulated datasets matched the true heterogeneity levels, following established thresholds: ^31,32^ none = 0.001, small = 0.1, medium = 0.25, and large ≥0.4. Detailed descriptions and contextual justification for these threshold values are provided in the supplementary information. By doing so, each simulation scenario represents a distinct and interpretable level of cross-chemical heterogeneity, enabling systematic evaluation of the hierarchical calibration framework’s performance across varying data structures.

The simulated data consisted of doses, responses of *KE*_*up*_ and responses of *KE*_*down*_ for each chemical. These datasets were fitted separately using both flat and hierarchical model specifications. The difference in predictive performance between the two specifications was evaluated using the differences in LOO and WAIC values. To ensure the stability of results across different response-response parameter values, the simulation procedure was repeated 100 times at each heterogeneity level, with performance metrics averaged across all iterations.

### 2.7 Case study: Non-mutagenic Liver Tumors Induced by Hepatic Proliferation

A case study was included to demonstrate the application of the proposed calibration approach. The case study utilizes multi-chemical rodent data from a previous study ^33^ on non-mutagenic tumorigenesis induced by sustained hepatic cell proliferation. These data are supported by a key event relationship “*Induction, Sustained Cell Proliferation leads to Formation, Liver tumor*” (Key Event Relationship ID 298^34^) in the AOP of “*Inhibition of iNOS, hepatotoxicity, and regenerative proliferation leading to liver tumors*” (AOP ID 32^35^). Multiple Doses per chemical were administered to male and female Fisher F344 and Sprague-Dawley rats. Sustained liver cell proliferation was quantified as the percentage of BrdU-positive cells using the 5-bromo-2-deoxyuridine (BrdU) incorporation assay. ^36^ Extra hepatic tumor incidence in rats was computed from tumor incidence after adjusting for background incidence. Detailed experimental protocols and quality control procedures are available in the original publication. ^33^ The complete dataset comprises two dose-response datasets for BrdU(%) (*KE*_*up*_) and tumor incidence (*KE*_*down*_) and is provided in the Supplementary Information. This dataset satisfies the essential requirements of the proposed calibration approach, including internal liver doses (ng/g), BrdU measurements, and tumor incidence values for each dose group and chemical.

The data were fitted using two models based on the qAOP framework (Equation (5)): one with a flat structure and one with a hierarchical structure. To isolate the impact of hierarchical calibration, both models employ identical dose-response structures, thereby controlling for uncertainty arising from dose-response modeling. Predictive performance was evaluated based on the models’ ability to predict both dose-response and response-response relationships, providing a comprehensive assessment of model accuracy across the entire pathway.

## 3 Results and Discussions

### 3.1 Results of Simulation Study

The response-response data generated through the iterative parameter adjustment procedure are visualized in Figure 2, illustrating how the true heterogeneity level determines parameter values and controls the shapes and locations of the simulated curves. In the absence of heterogeneity, parameters were concentrated around single point values to achieve Cohen’s f = 0.05, resulting in nearly identical response-response curves across all chemicals. At the small heterogeneity level, parameter values exhibited modest dispersion, yielding simulated data with Cohen’s f = 0.1. This level produced homogeneous curve shapes across chemicals with only slight variations in location. When heterogeneity increased to the medium level (Cohen’s f = 0.2), both the shapes and locations of the curves began to diverge noticeably across chemicals. At the large heterogeneity level, the simulated data achieved Cohen’s f = 0.5, with response-response curves exhibiting substantial variation in shape, location, and range across chemicals. These systematic differences demonstrate that the simulation procedure successfully generates datasets spanning a wide spectrum of cross-chemical heterogeneity, enabling robust evaluation of the hierarchical calibration framework.

**Figure 2.**
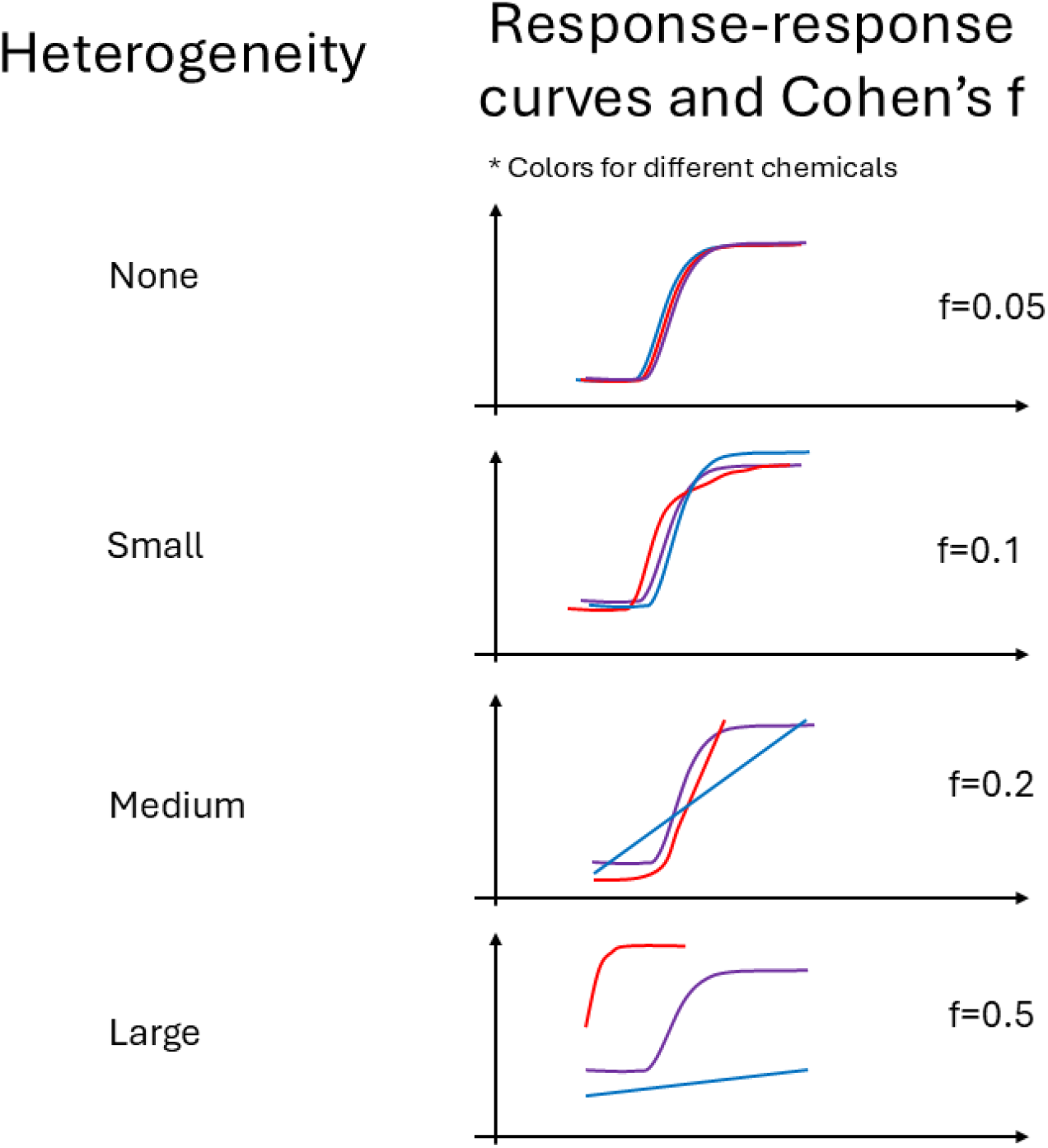
Diagram: Heterogeneity Propagates to Cohen’s f across Chemicals

After fitting both models to the simulated datasets, the model performance metrics (WAIC and LOO) demonstrated that the advantage of the hierarchical model over the flat model increases with the level of heterogeneity (Table 2). The comparative analysis revealed several key patterns. First, when heterogeneity exceeds the small level, both metrics consistently favor the hierarchical model. Under these conditions, the flat model conflates chemical-agnostic pathway effects with heterogeneity-induced variance, compromising predictive accuracy. Conversely, when heterogeneity is minimal (none or small), the flat model may perform comparably or even slightly better due to its reduced parameter complexity. However, as heterogeneity increases, the flat model’s inability to separate cross-chemical variation from underlying pathway effects leads to substantially degraded performance. Second, WAIC and LOO exhibited modestly divergent preferences for the hierarchical model at the medium heterogeneity level. Although both metrics are appropriate for Bayesian model comparison, LOO is generally preferred over WAIC ^25^ because they handle predictive performance evaluation and penalization of over-parameterization differently. In this study, the use of non-informative priors rendered the posterior distributions largely data-driven, with a limited number of extreme observations. As WAIC relies on variance-based approximations computed from the full dataset, it is more susceptible to the influence of such extremes. In contrast, LOO evaluates predictive performance by assessing changes in the posterior when individual observations are omitted, yielding a more robust basis for model comparison under these conditions. Overall, the performance differential between the flat and hierarchical models accurately reflects the underlying level of cross-chemical heterogeneity in the simulated data. These results provide strong empirical support for the validity of the proposed hierarchical calibration methodology.

**Table 1.**
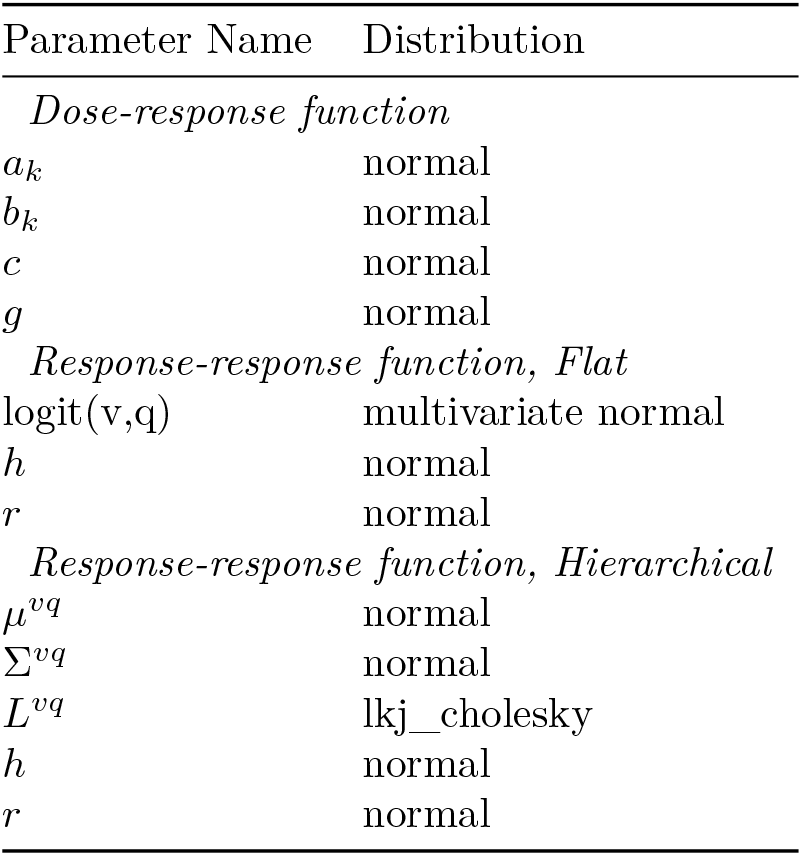
Prior Distributions for the parameters of the chosen models in the statistical calibration.

**Table 2.** Comparison of Predictive Performances between Flat and Hierarchical Models Across Different Heterogeneity Levels.

| Hetero | Cohen’s f | WAIC |  | LOO |  |  |
| --- | --- | --- | --- | --- | --- | --- |
|  |  | Diff<sup>1</sup> | P(Hier)<sup>2</sup> | Diff<sup>1</sup> | SE | P(Hier)<sup>2</sup> |
| 0=None | 0.05 | -0.52(1.83) | 3% | -5.70(2.66) | 1.85(0.69) | 8% |
| 1=Small | 0.1 | -0.06(3.68) | 42% | -0.99(1.96) | 1.75(0.36) | 44% |
| 2=Medium | 0.2 | 1.22(1.82) | 66% | 5.51(3.09) | 2.15(0.76) | 94% |
| 3=Large | 0.5 | 9.34(2.73) | 100% | 12.3(4.01) | 4.04(1.29) | 100% |

<sup>1</sup> mean and standard deviation (in parenthesis) over iterations, positive means in support of hierarchical model
<sup>2</sup> proportion of iterations supporting hierarchical model

### 3.2 Results of Case Study

For the case study using real data, the true heterogeneity level is unknown. The response-response data are predominantly concentrated in the low (log-BrdU ≤0.5, 71%) and medium (0.5-2, 24%) *KE*_*up*_ ranges (points in Figure 3), with limited information available in the high *KE*_*up*_ range (log-BrdU ≥2, 5%) from chemical 3,4,11. The spatial distribution of the response-response data suggests small to medium heterogeneity. The majority of data in the low *KE*_*up*_ range exhibit minimal variation across chemicals, although two data points from chemicals 7 and 9 fall outside the primary concentration area. In the high *KE*_*up*_ range, two points from chemicals 3 and 11 display log-BrdU values (~0.25) substantially lower than other points in this range (~0.8–0.9). These observations are consistent with the empirical Cohen’s f value of 0.17, indicating a small to medium ratio of between-to within-chemical variance. Based on the simulation study results, a hierarchical model is expected to demonstrate equivalent or marginally superior performance for data with small to medium heterogeneity.

**Figure 3.**
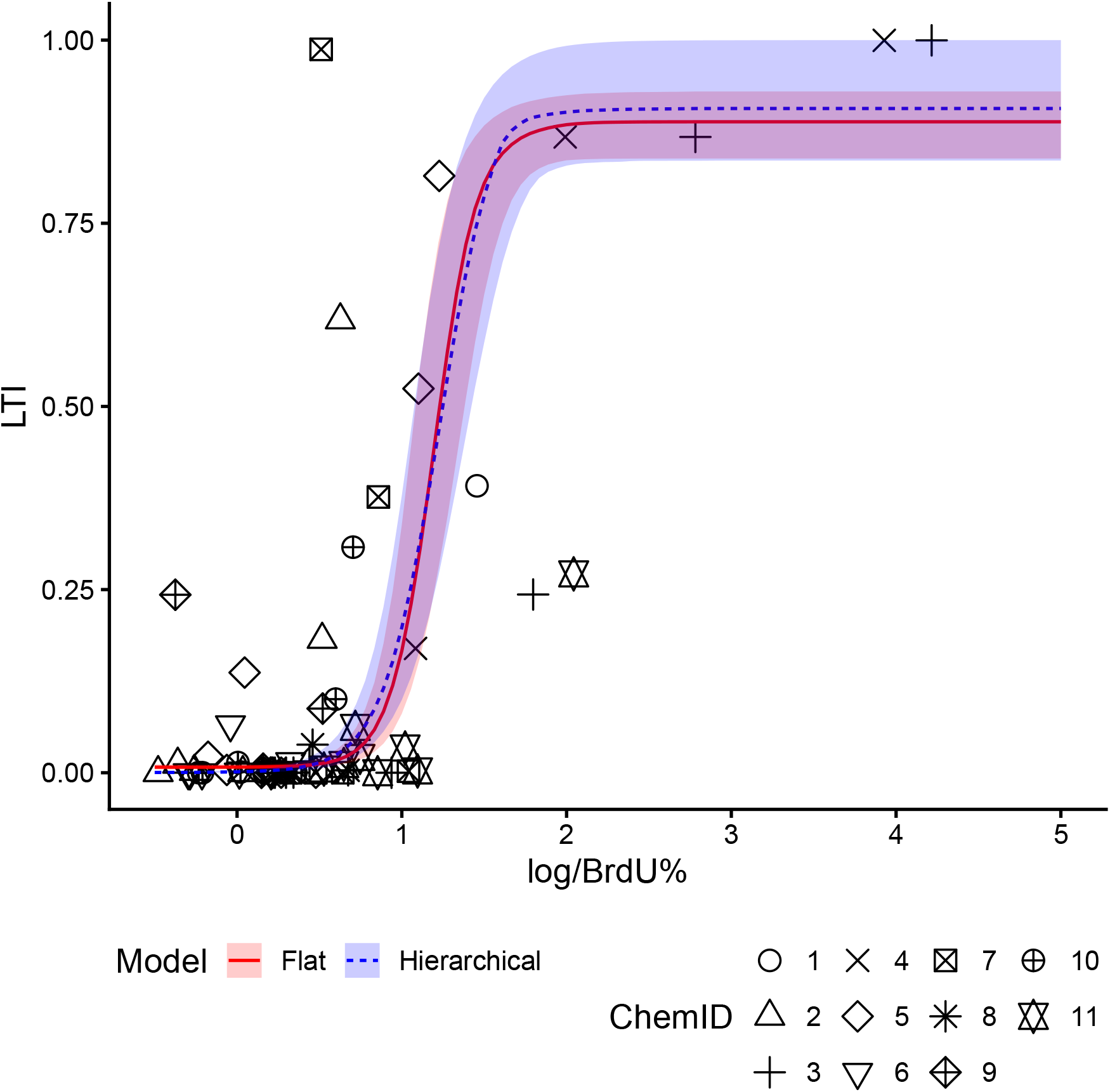
Comparing Flat and Hierarchical Response-response Models for Predicting Liver Tumor Incidence from Log-BrdU%. LTI: liver tumor incidence. Ribbons are 95% probability intervals

This expectation was confirmed by the model comparison results, which modestly favor the hierarchical model (WAIC_diff = 13.2(6.1) and LOO_diff = 18.2(6.7). This indicates that cross-chemical differences in the case study data are considerable and that the flat model failed to adequately capture this heterogeneity. Consequently, the flat model cannot be considered truly chemical-agnostic. In contrast, the hierarchical model successfully separates chemical-specific random effects from chemical-agnostic pathway effects, with the magnitude of random effects for each chemical visualized in the Supplementary Information.

A detailed graphical examination of the fitting results reveals additional insights. Both models fit the data well in the low *KE*_*up*_ range, where data concentration is highest. The predicted curves from the two models are nearly indistinguishable in this range, with the hierarchical model curve (blue dotted line) positioned slightly left of the flat model curve (red solid line). However, differences emerge in the high *KE*_*up*_ range. Although the mean curves remain similar, the 95% probability intervals of the hierarchical model (blue ribbon) provide substantially better coverage of the data compared to the flat model (red ribbon). Specifically, the hierarchical model exhibits broader upper and lower bounds that accommodate extreme data points from chemicals 3, 4, and 7. This reflects the higher dispersion estimated by the hierarchical model in the maximum probability, potency, and power parameters (v, h, and p, respectively).

Model goodness-of-fit could be influenced by both model structure and prior specifications. Since the dose-response components of both models were identical and prior distributions were the same, all observed performance differences are attributable to the response-response model structure. Therefore, the superior predictive performance of the hierarchical model results directly from the hierarchical structures implemented through the proposed calibration approach. Overall, these results demonstrate that the heterogeneity in the case study data is sufficiently substantial that the flat model violates the chemical-agnostic principle, whereas the hierarchical framework provides a valid methodological approach for identifying and characterizing cross-chemical heterogeneity.

### 3.3 Demonstrating Application with Case Study Results

To demonstrate the pragmatic application of the chemical-agnostic calibrated qAOP, we computed benchmark dose values for *KE*_*up*_ with a benchmark response (BMR) of extra 5% increase in tumor incidence, following Equation (9) from U.S. EPA ^18^ and EFSA guidance. ^19^

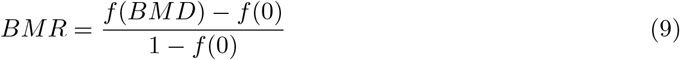

BMD values were calculated from the estimated model parameters as the doses corresponding to a pre-determined benchmark response (BMR) level. For the hierarchical model, a BMD distribution was computed using only the chemical-agnostic parameters. The derived values are hereafter referred to as benchmark “levels” (BMLs) rather than “doses” to emphasize that they are derived from BrdU rather than internal liver doses in a response-response qAOP framework.

The mean, lower bound, and upper bound of the computed BMLs are presented in Table 3 as BML, BMLL, and BMLU, respectively. The table also includes the ratio of mean to lower bound values (BML/BMLL), which provides two important insights.

**Table 3.** Benchmark Levels of BrdU% (regular and log scale) on Tumor Incidence Comparing Flat and Hierarchical Model Estimates.

| Model |  | Flat | Hierarchical |
| --- | --- | --- | --- |
| BrdU | BML | 1.32 | 0.67 |
|  | BMLL | 1.05 | 0.62 |
|  | BMLU | 1.61 | 0.99 |
| Ln(BrdU)<sup>1</sup> | BML | 0.121 | -0.174 |
|  | BMLL | 0.021 | -0.208 |
|  | BMLU | 0.207 | -0.004 |
| **BML/BMLL** |  | **1.25** | **1.08** |

<sup>1</sup> Same scale as x axis in Figure 3

First, the hierarchical model yields a lower BML/BMLL ratio (1.08) compared to the flat model (1.25), corroborating the previously reported performance metrics (WAIC_diff and LOO_diff). The BML/BMLL ratio reflects the level of uncertainty between mean and lower bound estimates, attributable to both data heterogeneity and model performance. A model is considered empirically reliable if this ratio falls within the range of 1 to 3, ^20^ and for model comparison purposes, lower ratios indicate superior performance. ^20^ Since both models are fitted to the same dataset, uncertainty arising from data heterogeneity is held constant. Consequently, the hierarchical model’s lower BML/BMLL ratio demonstrates superior characterization of uncertainty compared to the flat model.

Second, the BML, BMLL, and BMLU values from the hierarchical model are consistently lower than those from the flat model at a 5% BMR. These differences remain below two-fold, consistent with the relatively small to medium heterogeneity among chemicals in the low *KE*_*up*_ range. This finding aligns with the graphical patterns shown in Figure 3, where the hierarchical model curve is positioned to the left of the flat model curve. From a regulatory perspective, the conservative choice for the point of departure in risk assessment ^19,20^ would be the lower BML (or BMLL) value derived from the hierarchical model. It is important to note that these differences in BML values should not be interpreted as direct indicators of model performance. According to EPA BMDS guidelines, a model yielding lower BMD values should not be recommended if its performance metrics (e.g., AIC in BMDS) are inferior to those of alternative candidate models. ^18^ In this case study, however, the hierarchical model demonstrates both superior performance metrics and lower, more conservative BML values. Overall, these results demonstrate that a qAOP calibrated using the chemical-agnostic hierarchical approach not only provides superior model performance but also derives BML values that support robust, probabilistic risk assessment.

## 4 Conclusions

Achieving truly chemical-agnostic adverse outcome pathway (AOP) and quantitative AOP (qAOP) frameworks requires explicit recognition and systematic treatment of cross-chemical variability that can confound pathway-level inferences. Without proper characterization and separation of chemical-specific effects, fundamental pathway relationships become obscured by heterogeneity-induced noise, thereby compromising the reliability of chemical-agnostic predictions.

The chemical-agnostic calibration approach proposed in this study addresses this challenge by leveraging hierarchical structures to systematically separate chemical-specific heterogeneity from underlying pathway effects. Through this methodological framework, chemical-specific deviations are explicitly modeled as random effects, enabling the extraction of pathway-level parameters that represent core mechanistic relationships independent of individual chemical properties.

The validity of this framework is supported by rigorous mathematical principles and demonstrated performance in a simulation study, which yield two key conclusions. First, performance differences between models with and without hierarchical calibration can be used to reveal the magnitude of heterogeneity in the data. This was shown when the hierarchical model received higher performance when heterogeneity reached at least medium levels. Second, when heterogeneity is substantial, an uncalibrated qAOP should not be considered truly chemical-agnostic in practice, as it confounds pathway-level effects with chemical-specific variation. Therefore, the hierarchical calibration approach proposed in this study represents a critical methodological advancement for developing robust chemical-agnostic AOP frameworks applicable to next-generation risk assessment.

## 5 Acknowledgement

This project received funding from the European Union’s Horizon 2020 Research and Innovation programme under Grant Agreement No. 964537 (RISK-HUNT3R), and it is part of the ASPIS cluster. The opinions expressed in this document reflect only the authors’ view. The European Commission is not responsible for any use that may be made of the information it contains.

We thank Veltman, Christina H.J. and Khalidi, Hiba and Zgheib, Elias and van de Water, Bob and Luijten, Mirjam and Pennings, Jeroen L.A. for sharing the data and support for this study.

## 6 Supporting Information

1. Supporting Information.pdf. Contents: 1) glossary of acronyms. 2) dose-response functions. 3) formula to compute Cohen’s f effect size. 4) prior specifications. 5) metadata codebook for the liver case study data. 6) figures of fitted chemical-specific qAOP models for the liver case study. 7) stan code for the benchmark dose models. 8) R code for the simulation study.
2. Data of case study from: ^33^ Veltman2025.xlsx

## Notes

### Competing Interest Statement

The authors have declared no competing interest.

### Summary of Updates

Revision submitted to Chemical Research in Toxicology

