## Supplementary Information for "A Method to Calibrate Chemical-agnostic Quantitative Adverse Outcome Pathways on Multiple Chemical Data"

#### Contents


**1 Glossary of Acronyms** **S2**

**2 Benchmark Dose Functions** **S2**

**3 Calcuating Cohen's f effect size** **S3**

**4 Prior specification** **S3**

**5 Data of case study** **S5**

**6 Fitted Curves of Chemical-specific Random Effets by the Hierarchical Model** **S8**

**7 Stan code for qAOPs** **S8**

**8 R script for simulation study** **S13**

### 1 Glossary of Acronyms

- AOP: adverse outcome pathway
- AO: adverse outcome
- qAOP: quantitative adverse outcome pathway
- KE: key event
- NAM: new approach methdology
- POD: point of departure
- MOA: mode of action
- BMD: benchmark dose
- BMDS: benchmark dose software
- LOO-CV: leave-one-out cross validation
- PSIS: pareto smoothed importance sampling
- ELPD: expected log predictive density
- AIC: Akaike information criteria
- WAIC: widely applicable information criteria
- MCMC: Markov chain Monte Carlo
- BrdU: 5 -bromo-2 -deoxyuridine
- PDF: probability density function
- BMR: benchmark response
- BML: benchmark level

### 2 Benchmark Dose Functions

Candidate functions for continuous and dichotomous types of responses were adapted from the U.S. EPA Benchmark Dose Software.[1]

Table S1: Continuous models and parameterizations

| Model | Parameterization | Additional Specifications |
|---|---|---|
| Linear | f(dose)=a+b*dose;<br>a = control response;<br>b= slope | b>0 when increasing, b<0 when decreasing |
| Power | f(dose)=a+b*dose^g;<br>a = control response;<br>b = slope;<br>g = power | 0<=g<=18;<br>g may be further restricted to g >= 1 to avoid infinite slope at the control dose. |
| Polynomial | f(dose)=a+b_1*dose+b_2*dose^2+...b_n*dose^n;<br>a = control response;<br>b_1,...b_n = polynomial coefficients;<br>n = degree of polynomial | The degree of polynomial should not exceed the number of dose group minus one |
| Hill | f(dose)=a+b*dose^g/(c^g+dose^g);<br>a = control response;<br>b = maximum change;<br>c = dose with half-maximal change;<br>g = power | It is recommended to normalize the response so that the half-maximal dose could be constrainted: 0<=c<=5<br>0<=g<=18, g may be further restricted to g>= 1 |
| Exponential | Exp2: f(dose)=a * exp(±(b*dose));<br>Exp3: f(dose)=a * exp(±(b*dose)^d);<br>Exp4: f(dose)=a*(c-(c-1)*exp(-b*dose));<br>Exp5: f(dose) = a * (c-(c-1)*exp(-(b*dose)^d));<br>a = control response;<br>b = slope;<br>c = asymptote term;<br>d = power | a>0;<br>Direction of change is controlled by the ± sign and parameter c;<br>0<b<100;<br>0<c<1 when decrasing; c>1 when increasing;<br>1<=d<=18 |

### 3 Calcuating Cohen's f effect size

$$\begin{aligned} ES &= \sqrt{\frac{SSB}{SST - SSB}} \\ SSB &= \sum_{j=1}^{K} n_j(\mu_j - \hat{\mu})^2 \\ SST &= \sum_{i=1}^{N} (y_i - \hat{\mu})^2 \end{aligned} \tag{1}$$

where SSB and SST are between-chemical and total squared variance, respectively; $\hat{\mu}$ is global mean of the data; j is chemical index, $n_j$ and $\mu_j$ are the size and mean per chemical. The value of Cohen's f can be interpreted as follows:[2] when f=0, there is no between-chemical differences; when f>0, f=0.1 ~ small differences, f=0.2 ~ medium differences, f>=0.4 ~ large differences. The effect size values are calculated using Analysis of Covariate (ANCOVA) as the ratio of between-group and within group variances, adjusted for covariates.[3] These are rules of thumb, developed to assist the interpretation of Cohen's f effect sizes from psychological studies.[3] The values in the rules have been validated through simulation studies and updated.[2,4] A large Cohen's f value indicates the between-group variance is bigger than that within groups.

### 4 Prior specification

Prior choices for the simulation study. $k$ is chemical index.

Dose-response part (shared by flat and hierarchical model):

Table S2: Dichotomous models and parameterizations

| **Model** | **Parameterization** | **Additional Notes** |
|---|---|---|
| Quantal-Linear | p(dose)=a+(1−a)×[1−exp(−b×dose)];<br>a = background;<br>b = slope | 0<=a<=1;<br>0<b<100; |
| Probit | p(dose)=Φ(a+b*dose);<br>a = intercept;<br>b = slope | b>=0 |
| Logistic | p(dose)=1/[1+exp(-a-b*dose)];<br>a = intercept;<br>b = slope | b>=0 |
| Multistage | p(dose)=a+(1-a)(1-exp[-\sum_{j=1}^n{b_j*dose^j}]);<br>a = background;<br>b_j = dose coefficient | j<=23;<br>0<=a<=1;<br>b_j can be stricted to b>=0 which guarantee that the model will be either flat or always increasing |
| Weibull | p(dose)=a+(1-a)*(1-exp[-b*dose^g]);<br>a = background;<br>b = slope;<br>g = power | 0<=a<=1;<br>0<b<100;<br>0<g<=18, g can be stricted to g>=1 |
| LogLogistic | p(dose)=a+((1-a)/{1+exp[-b-g*log(d)]});<br>a = background;<br>b = slope;<br>g = power | 0<=a<=1;<br>0<g<18, g could be restricted to g>=1 |
| LogProbit | p(dose)=a+(1-a)*Φ[b+g*log(d)];<br>a = background;<br>b = slope;<br>g = power | 0<=a<=1;<br>0<g<18, g could be restricted to g>=1 |
| Dichotomous Hill | p(dose)=v*(1+r*exp(-(h+q*log(x))))/(1+r*exp(-(h+q*log(x))));<br>v = maximum probability;<br>v*q = background probability, q = extra risk;<br>h = potency;<br>r = power | 0<v<=1;<br>0<=q<1;<br>-18<h<=18;<br>0<=p<=18, can be restricted to p>=1; |

$$
\begin{aligned}
\log(Y1) &= a_k + \frac{b_k \cdot dose^g}{c^g + dose^g} \\
a_k &\sim normal(0,1) \\
b_k &\sim normal(0,1) \\
c &\sim normal(0,5) \\
g &\sim normal(0,1)
\end{aligned}
\tag{2}
$$

Response-response part, flat model:

$$
\begin{aligned}
p(Y2=1) &= v + \frac{1 + q \cdot \exp(-(h + r \cdot \log(Y1)))}{1 + \exp(-(h + r \cdot \log(Y1))} \\
logit(v,q) &\sim MVN([0,0]', [1,1]') \\
h &\sim normal(0,1) \\
r &\sim normal(0,1)
\end{aligned}
\tag{3}
$$

Response-response part, hierarchical model:

$$
\begin{aligned}
p(Y2=1) &= v_k + \frac{1 + q_k \cdot \exp(-(h + r \cdot \log(Y1)))}{1 + \exp(-(h + r \cdot \log(Y1))} \\
v_k &= \mu^v + \sigma^v + z_k^v \\
q_k &= \mu^q + \sigma^q + z_k^q \\
logit(v, q) &\sim MVN(\mu^{vq}, \Sigma^{vq} \cdot L^{vq}) \\
\mu^{vq} &= (\mu^v, \mu^q) \\
\Sigma^{vq} &= (\sigma^v, \sigma^q) \\
L_k^{vq} &= (z_k^v, z_k^q) \\
\mu^{vq} &\sim normal(0, 1) \\
\Sigma^{vq} &\sim normal(0, 5) \\
L^{vq} &\sim lkj\_cholesky(2) \\
h &\sim normal(0, 1) \\
r &\sim normal(0, 1)
\end{aligned} \tag{4}
$$

### 5 Data of case study

From the case study of non-mutagenic tumorgenesis induced by sustained cell proliferation in liver.[5] The data is provided as Veltman2025.csv and includes the following variables:

1. N_subj: number of animals per dose group in BrdU measurements
2. Unit: reported unit of dose in the source report
3. Dose: doses from the source report, of ‘Route‘ at ‘Unit‘.
4. Unit_liver: unit of converted internal liver doses from[5]
5. Liver: converted internal liver doses from[5]
6. Mod: modifying factor to unify internal liver doses to µg/mL
7. Liver_M: unified internal liver doses (µg/mL) = ‘Liver‘ * ‘Mod‘
8. BrdU_mean: $KE_{up}$ BrdU percentage, group mean
9. BRdU_SD: $KE_{up}$ BrdU percentage, group standard deviation
10. N_tumour: number of animals per dose group in liver tumor measurements
11. Extra_tumour: extra liver tumor incidence per dose group

The dose-response data is illustrated as Figure S1.

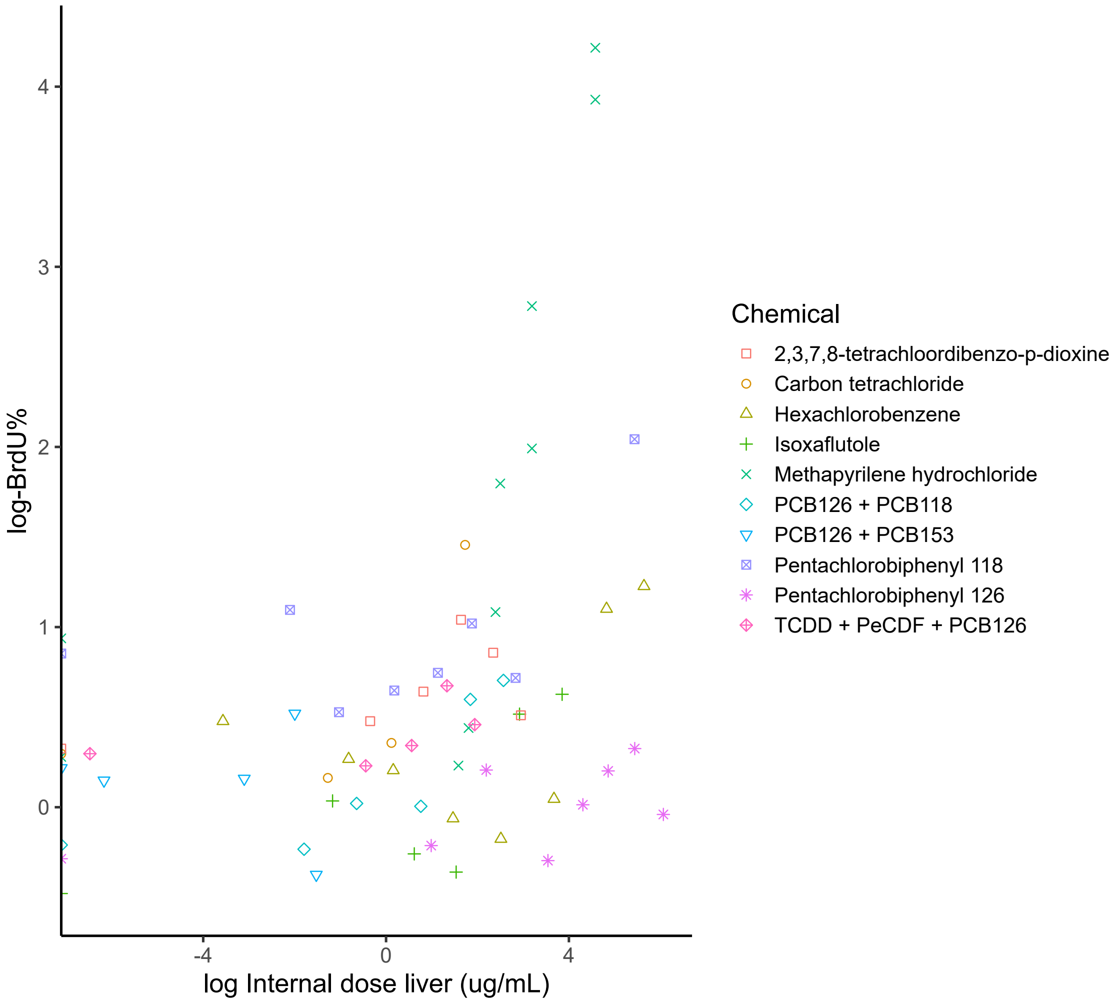


Figure S1: Dose-response Scatter Plot. Source: Veltman et al. (2025)

The fitted chemical-specific dose-response and response-response random effects are plotted as Figure S2 and S3 respectively.

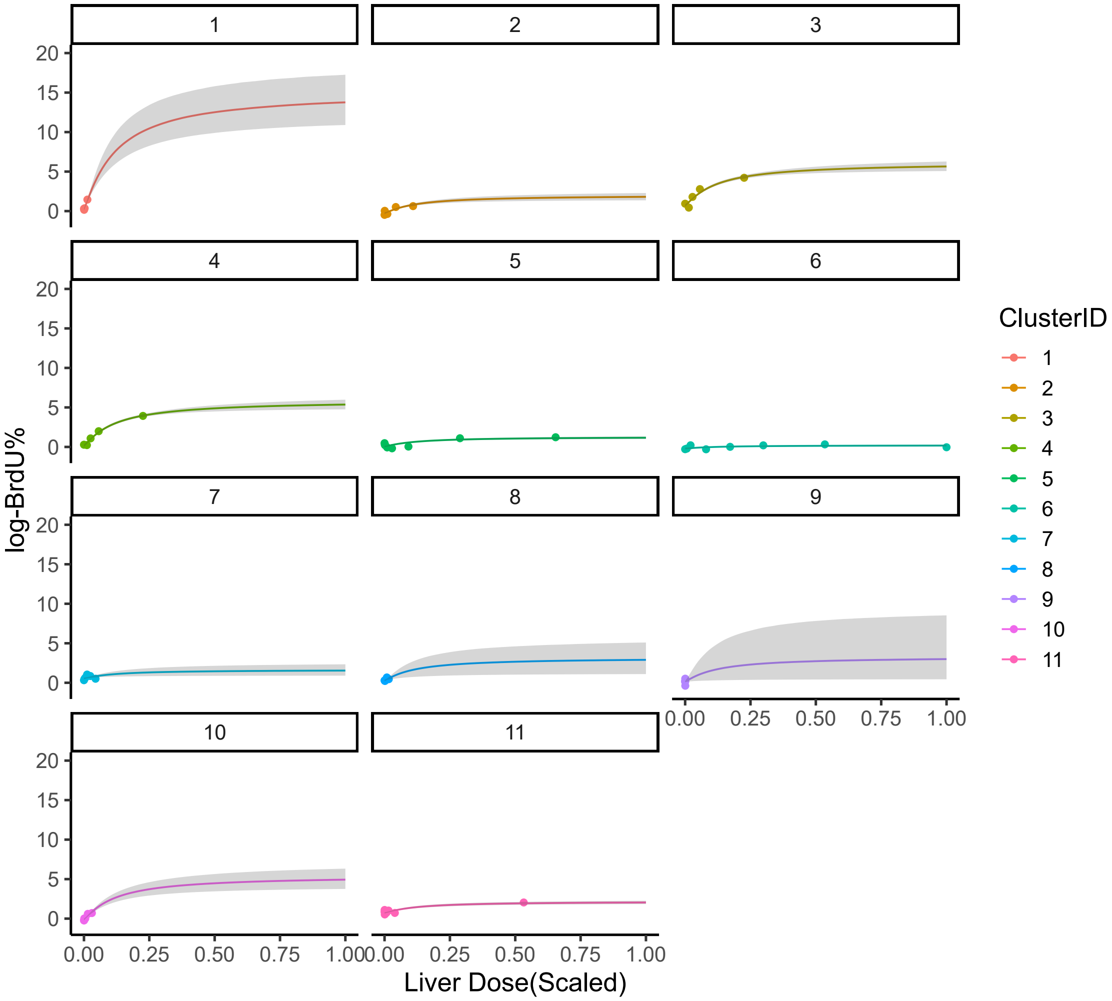


Figure S2: Chemical-specific Dose-response Curves of Random Effects in Sustained Liver Cell Proliferation Induced by Chemical Exposure in Rodents

### 6 Fitted Curves of Chemical-specific Random Effets by the Hierarchical Model

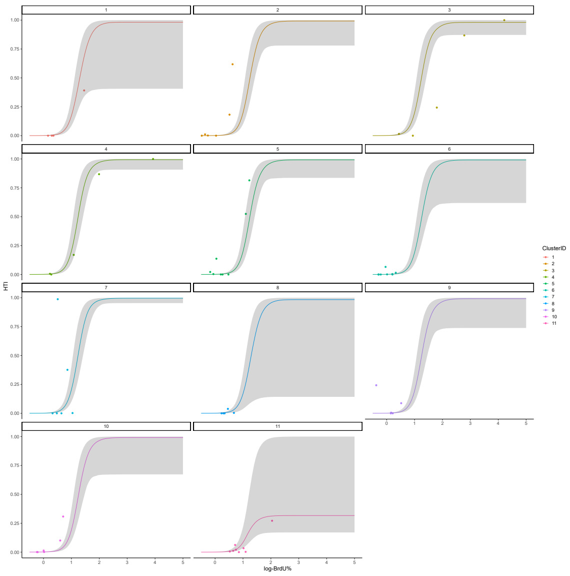


Figure S3: Chemical-specific Response-response Random Effects of Liver Tumor Incidence Predicted by Log-BrdU% by the Hierarchical Model

### 7 Stan code for qAOPs

Flat model

```
functions{
  // Updated density function for sigma
  real Normal_Summary_lpdf(array[] real ymean, array[] real ysd, array
      [] int Nsub,
    // array[] real fd, array[] real sigma){
    // return -sum(to_vector(Nsub) .* log(to_vector(sigma))+ ((
        to_vector(Nsub)-1).*square(to_vector(ysd))+(to_vector(Nsub)).*
        square(to_vector(ymean)-to_vector(fd)))./(2*to_vector(sigma)^2))
        ;
    array[] real fd, real sigma){
    return -sum(to_vector(Nsub) .* log((sigma))+ ((to_vector(Nsub)-1).*
        square(to_vector(ysd))+(to_vector(Nsub)).*square(to_vector(ymean
        )-to_vector(fd)))./(2*(sigma)^2));
  }
  array[] real pow_vec(array[] real x, real power){
    array[size(x)] real x_power;
```

```
    for(n in 1:size(x)){
      x_power[n] = pow(x[n], power);
    }
    return x_power;
  }
}
data{
  int run_simu;                          // simulate or estimate
  int<lower=1> S;                        // number of chemicals
  // Y1 data
  int N1;                                // number of Y1 groups
  array[N1] real<lower=0, upper=1> d1;  // dose for Y1 in (0, 1)
  array[N1] real Lymean1;                // log-mean for Y1
  array[N1] real<lower=0> Lysd1;                  // log-SD for Y1
  array[N1] int Nsub1;                   // subjects per Y1 group
  array[N1] int<lower=1, upper=S> ss1;  // chemical index for Y1
  // Y2 data
  int N2;                                // number of Y2 groups
  array[N2] real<lower=0, upper=1> d2;  // dose for Y2 in (0, 1)
  array[N2] int ycount2;                 // incidence count for Y2
  array[N2] int Nsub2;                   // subjects per Y2 group
  array[N2] int<lower=1, upper=S> ss2;  // chemical index for Y2
  // hyperpriors
  array[2] real prior_sigma;          // prior for sigma_Y1[s]
  array[2] real power_bound;             // lower and upper bounds for
     power g
  array[2] real prior_threshold;       // prior for threshold parameter
  array[2] real prior_background;      // prior for background parameter
  array[2] real prior_slope;          // prior for slope parameter
}
parameters{
  vector[S] a1;                          // chemical-specific intercept for Y1
  vector<lower=0>[S] b1;                 // chemical-specific slope for Y1
  real<lower=0> c1;                      // Hill function parameter for Y1 (
     not chemical-specific)
  real<lower=power_bound[1],upper=power_bound[2]> g1;// Hill function
     parameter for Y1 (not chemical-specific)
  real<lower=0> sigma_Y1;               // global sigma for Y1
  real<lower=0, upper=1> v2;            // background p for Y2
  real<lower=0, upper=1> b2;            // maximum p for Y2
  real a2;                               // potency for Y2
  real<lower=power_bound[1],upper=power_bound[2]> g2;// power for Y2
  vector[N2] z_Y2;                       // latent errors for Y2
}
transformed parameters{
  real z_v2 = logit(v2);
```

```
  real z_b2 = logit(b2);
  array[N1] real Lmu1d1;          // log-Y1 mean at d1 (per chemical,
     Hill function)
  array[N2] real Lmu1d2;          // latent log-Y1 mean at d2 (per
     chemical, Hill function)
  array[N2] real<lower=0> y1d2;// latent Y1 at d2 in (0, 1)
  array[N2] real<lower=0, upper=1> p2d2; // latent probability for Y2
  // Y1 mean at d1 (chemical-specific, Hill function)
  for(n1 in 1:N1){
    Lmu1d1[n1] = (a1[ss1[n1]] + (b1[ss1[n1]] * pow(d1[n1], g1)) / (pow(
       c1, g1) + pow(d1[n1], g1)));
  }
  // Y1 mean at d2 and link to Y2 (chemical-specific, Hill function)
  for(n2 in 1:N2){
    Lmu1d2[n2] = (a1[ss2[n2]] + (b1[ss2[n2]] * pow(d2[n2], g1)) / (pow(
       c1, g1) + pow(d2[n2], g1)));
    y1d2[n2] = exp(Lmu1d2[n2] + z_Y2[n2] * sigma_Y1);
    p2d2[n2] = v2 * (1 + b2 * exp(-(a2 + g2 * log(y1d2[n2])))) /
               (1 + exp(-(a2 + g2 * log(y1d2[n2]))));
  }
}
model{
  // priors
  sigma_Y1 ~ cauchy(prior_sigma[1], prior_sigma[2]);
  a1 ~ normal(prior_threshold[1], prior_threshold[2]);
  b1 ~ normal(prior_threshold[1], prior_threshold[2]);
  c1 ~ normal(prior_background[1], prior_background[2]);
  g1 ~ normal(prior_threshold[1], prior_threshold[2]);
  v2 ~ beta(prior_slope[1],prior_slope[2]);
  b2 ~ beta(prior_slope[2],prior_slope[1]);
  vector[2] z_vb;
  z_vb[1] = z_v2;
  z_vb[2] = z_b2;
  z_vb ~ multi_normal([0,0]',diag_matrix([1,1]'));
  // v2 ~ beta(prior_slope[1], prior_slope[2]);
  // b2 ~ beta(prior_slope[1], prior_slope[2]);
  a2 ~ normal(prior_threshold[1], prior_threshold[2]);
  g2 ~ normal(prior_threshold[1], prior_threshold[2]);
  z_Y2 ~ std_normal();
  // Model
  if(run_simu == 0){
    target += Normal_Summary_lpdf(Lymean1 | Lysd1, Nsub1, Lmu1d1,
       sigma_Y1);
    ycount2 ~ binomial(Nsub2, p2d2);
  }
}
```

Hierarchical model

```
functions{
  // Updated density function for per-chemical sigma
  real Normal_Summary_lpdf(array[] real ymean, array[] real ysd, array
      [] int Nsub,
    // array[] real fd, array[] real sigma){
    // return -sum(to_vector(Nsub) .* log(to_vector(sigma))+ ((
        to_vector(Nsub)-1).*square(to_vector(ysd))+(to_vector(Nsub)).*
        square(to_vector(ymean)-to_vector(fd)))./(2*to_vector(sigma)^2))
        ;
    array[] real fd, real sigma){
    return -sum(to_vector(Nsub) .* log((sigma))+ ((to_vector(Nsub)-1).*
        square(to_vector(ysd))+(to_vector(Nsub)).*square(to_vector(ymean
        )-to_vector(fd)))./(2*(sigma)^2));
  }
  array[] real pow_vec(array[] real x, real power){
    array[size(x)] real x_power;
    for(n in 1:size(x)){
      x_power[n] = pow(x[n], power);
    }
    return x_power;
  }
}
data{
  int run_simu;                     // simulate or estimate
  int<lower=1> S;                   // number of chemicals
  // Y1 data
  int N1;                           // number of Y1 groups
  array[N1] real<lower=0, upper=1> d1;  // dose for Y1 in (0, 1)
  array[N1] real Lymean1;               // log-mean for Y1
  array[N1] real<lower=0> Lysd1;                 // log-SD for Y1
  array[N1] int Nsub1;                  // subjects per Y1 group
  array[N1] int<lower=1, upper=S> ss1;  // chemical index for Y1
  // Y2 data
  int N2;                           // number of Y2 groups
  array[N2] real<lower=0, upper=1> d2;  // dose for Y2 in (0, 1)
  array[N2] int ycount2;                // incidence count for Y2
  array[N2] int Nsub2;                  // subjects per Y2 group
  array[N2] int<lower=1, upper=S> ss2;  // chemical index for Y2
  // hyperpriors
  array[2] real prior_sigma;          // prior for sigma_Y1[s]
  array[2] real power_bound;              // lower and upper bounds for
     power g
  array[2] real prior_threshold;       // prior for threshold parameter
  array[2] real prior_background;      // prior for background parameter
  array[2] real prior_slope;          // prior for slope parameter
```

```
}
parameters{
  // dose-response
  vector[S] a1;                    // chemical-specific intercept for Y1
  vector<lower=0>[S] b1;           // chemical-specific slope for Y1
  real<lower=0> c1;                // Hill function parameter for Y1 (
      not chemical-specific)
  real<lower=power_bound[1],upper=power_bound[2]> g1;// Hill function
      parameter for Y1 (not chemical-specific)
  real<lower=0> sigma_Y1;          // global sigma for Y1
  // response-response
  vector[2] mu_vb;                 // Mean vector for logit(v2) and
      logit(b2)
  cholesky_factor_corr[2] L_vb;    // Cholesky decomposition of
      correlation matrix
  vector<lower=0>[2] sigma_vb;     // Standard deviations for logit(v2)
      and logit(b2)
  matrix[2, S] z_raw_vb;           // Raw latent deviations for v2 and
      b2
  real a2;                         // global
  real<lower=power_bound[1],upper=power_bound[2]> g2; // global
  vector[N2] z_Y2;                 // latent errors for Y2
}
transformed parameters{
  vector<lower=0,upper=1>[S] v2;
  vector<lower=0,upper=1>[S] b2;
  for (s in 1:S) {
    vector[2] logit_vb = mu_vb + diag_pre_multiply(sigma_vb, L_vb) *
        z_raw_vb[:, s];
    v2[s] = inv_logit(logit_vb[1]);
    b2[s] = inv_logit(logit_vb[2]);
  }
  array[N1] real Lmu1d1;           // log-Y1 mean at d1 (per chemical,
      Hill function)
  array[N2] real Lmu1d2;           // latent log-Y1 mean at d2 (per
      chemical, Hill function)
  array[N2] real<lower=0> y1d2;             // latent Y1 at d2 in (0, 1)
  array[N2] real<lower=0, upper=1> p2d2; // latent probability for Y2
  // Y1 mean at d1 (chemical-specific, Hill function)
  for(n1 in 1:N1){
    Lmu1d1[n1] = (a1[ss1[n1]] + (b1[ss1[n1]] * pow(d1[n1], g1)) / (pow(
        c1, g1) + pow(d1[n1], g1)));
  }
  // Y1 mean at d2 and link to Y2 (chemical-specific, Hill function)
  for(n2 in 1:N2){
    Lmu1d2[n2] = (a1[ss2[n2]] + (b1[ss2[n2]] * pow(d2[n2], g1)) / (pow(
```

```
        c1, g1) + pow(d2[n2], g1)));
    p2d2[n2] = v2[ss2[n2]] * (1 + b2[ss2[n2]] * exp(-(a2 + g2 * log(
        y1d2[n2])))) /
                (1 + exp(-(a2 + g2 * log(y1d2[n2]))));
  }
}
model{
  // priors
  sigma_Y1 ~ cauchy(prior_sigma[1], prior_sigma[2]);
  a1 ~ normal(prior_threshold[1], prior_threshold[2]);
  b1 ~ normal(prior_threshold[1], prior_threshold[2]);
  c1 ~ normal(prior_background[1], prior_background[2]);
  g1 ~ normal(prior_threshold[1], prior_threshold[2]);
  mu_vb ~ normal(prior_threshold[1], prior_threshold[2]);
  sigma_vb ~ cauchy(prior_sigma[1], prior_sigma[2]);
  L_vb ~ lkj_corr_cholesky(2);
  to_vector(z_raw_vb) ~ std_normal();
  a2 ~ normal(prior_threshold[1], prior_threshold[2]);
  g2 ~ normal(prior_threshold[1], prior_threshold[2]);
  z_Y2 ~ std_normal();
  // Model
  if(run_simu == 0){
    target += Normal_Summary_lpdf(Lymean1 | Lysd1, Nsub1, Lmu1d1,
        sigma_Y1);
    ycount2 ~ binomial(Nsub2, p2d2);
  }
}
```

### 8 R script for simulation study

Data was simulated to include ten chemicals. Four dose levels were randomly sampled per chemical within the range of 0 to 1 $\mu mol$ for $KE_{up}$ and $KE_{down}$ as $d_1$ and $d_2$, respectively. Each dose group included of ten subjects. The flat and hierarchical models share the same dose-response part, which was specified with chemical-specific hierarchical structures.

```
run_simu <- function(S,Hetero){
  c1 <- 0.5       # Half-maximal dose for CHill
  g1 <- 6.0       # Steep Hill coefficient for CHill (creates threshold)
  a2 <- -6      # Threshold location parameter in DHill
  g2 <- 3     # Steep DHill coefficient (creates threshold)

  # Parameters adjusted for different heterogeneity levels
  if(Hetero == 1) { # None, Cohen's f   0.01
    sigma1 = rep(0.01, S)
    a1 = rep(1, S)
    b1 = rep(2, S)
```

```
    v2_m = rbeta(S,50,1)
    b2_m = rbeta(S,1,50)
  } else if(Hetero == 2) { # Low, Cohen's f   0.1
    sigma1 = rnorm(S,0.3, 0.05)
    a1 = rnorm(S, 1, 0.1)
    b1 = rnorm(S, 2, 0.1)
    v2_m = rbeta(S,50,3)
    b2_m = rbeta(S,3,50)
  } else if(Hetero == 3) { # Cohen's f   0.2
    sigma1 = rnorm(S,0.6, 0.1)
    a1 = rnorm(S, 1, 0.25)
    b1 = rnorm(S, 2, 0.2)
    v2_m = rbeta(S,10,1)
    b2_m = rbeta(S,1,10)
  } else if(Hetero == 4) { # Cohen's f >= 0.4
    sigma1 = rnorm(S,0.8, 0.2)
    a1 = rnorm(S, 1, 0.3)
    b1 = rnorm(S, 2, 0.5)
    v2_m = rbeta(S,3,1)
    b2_m = rbeta(S,1,3)
  }
  # G1 <- runif(S,4,6) %>% round                ## number of dose groups
      per cluster
  G1 <- rep(6,S) %>% as.numeric()
  N1 <- sum(G1)                                 ## number of obs per
      cluster
  n1_m <- sapply(G1,function(g){                ## number of subjects per
      dose group
    n <- as.integer(10 + runif(g,0,0) )
    return(list(n))
  })
  d1_m <- sapply(G1,function(g){               ## dose per group
    d <- seq(0,1,length.out=g)
    return(list(d))
  })
  # G2 <- runif(S,4,6) %>% round
  G2 <- rep(6,S) %>% as.numeric()
  N2 <- sum(G2)
  d2_m <- sapply(G2,function(g){               ## dose per group
    return(list(d))
  })
  n2_m <- sapply(G2,function(g){
    n <- as.integer(10 + runif(g,0,0) )
    return(list(n))
  })
```

```
mean1_m <- lapply(as.list(1:S),function(s){        # compute E(y) per
    dose group
  fd <- get_CHill(d1_m[[s]],a1[[s]],b1[[s]],c1,g1) %>%
    as_tibble %>% add_column(d=d1_m[[s]])
  return(fd)
})

y1_ind_m <- lapply(as.list(1:S),function(s){     # simulate y
  gen <- gen_ind_con(n1_m[[s]],d1_m[[s]],mean1_m[[s]]$value,sigma1[s
      ],
                    lognormal = T,a=0,b=100)
  gen <- gen %>% mutate(ss = s)
  return(gen)
})
y1_sum_m <- lapply(as.list(1:S),function(s){    # summarize y by mean
    and sd
  sum <- gen_sum_con(y1_ind_m[[s]])
  sum %>% as_tibble %>%
    mutate(ss = s)
})
dir_b2 <- c(0,1)                          # max extra risk, vb is
    background probability
miny1 <- do.call("rbind",y1_ind_m)$y %>% min
maxy1 <- do.call("rbind",y1_ind_m)$y %>% max
m <- 1/N1 * 0.5
y2d2_ind_m <- lapply(as.list(1:S),function(s){
  pd2 <- get_DHill(y1_ind_m[[s]]$y,v=v2_m[[s]],b=b2_m[[s]],a=a2,g=g2)
  y2d2 <- y1_ind_m[[s]] %>%
    mutate(
      p = pd2
    ) %>% rename(y1=y) %>% rowwise() %>% mutate(
      y = rbern(1,p))
  gen <- y2d2 %>% mutate(
    ss = s
  )
  return(gen)
})
y2_sum_m <- lapply(as.list(1:S),function(s){    # summarize y by mean
    and sd
  sum <- gen_sum_dic(y2d2_ind_m[[s]])
  sum <- sum %>% as_tibble %>%
    mutate(
      ss = s
    )
})
df_ind_y1 <- do.call("rbind",y1_ind_m)
```

```
  df_ind_y2 <- do.call("rbind",y2d2_ind_m)
  df_sum_y1 <- do.call("rbind",y1_sum_m)
  df_sum_y2 <- do.call("rbind",y2_sum_m)
  return(list(df_ind_y1,df_ind_y2,df_sum_y1,df_sum_y2))
}

# Run simulation———
Hetero <- 1:4
S <- 10
iter <- 100
for(i in 1:iter){
  iterID <- (Hetero-1)*iter+i
  ls_simu <- run_simu(S=S,Hetero = Hetero)
  df_ind_y1 <- ls_simu[[1]]
  write.csv(df_ind_y1,
            file = file.path(dir_storage,paste0("T",iterID,"
              _y_ind_DR1_Hill.csv")),
            row.names = F)
  df_ind_y2 <- ls_simu[[2]]
  write.csv(df_ind_y2,file = file.path(dir_storage,paste0("T",iterID,"
     _y_ind_DR2_DHill.csv")),
            row.names = F)
  # get_ANCOVA(df_ind_y2$ss,df_ind_y2$p,df_ind_y2$dose)
  df_sum_y1 <- ls_simu[[3]]
  write.csv(df_sum_y1,file = file.path(dir_storage,paste0("T",iterID,"
     _y_sum_DR1_Hill.csv")),
            row.names = F)
  df_sum_y2 <- ls_simu[[4]]
  write.csv(df_sum_y2,file = file.path(dir_storage,paste0("T",iterID,"
     _y_sum_DR2_DHill.csv")),
            row.names = F)
}

# Test———
i_warmup <- 1e3
i_sample <- 1e3

datalist_ind <- data_to_stan_RR_ind_con(df_ind_y1,df_ind_y2)
# Ship to stan————
df_logy1 <- lnorm_logpars(df_sum_y1$ymean,df_sum_y1$ysd)
datalist_sum <- list(
      S = n_distinct(df_sum_y1$ss),
      N1 = nrow(df_sum_y1),
      N2 = nrow(df_sum_y2),
      ss1 = df_sum_y1$ss %>% as.integer(),
      ss2 = df_sum_y2$ss %>% as.integer(),
```

```
        Nsub1 = df_sum_y1$n %>% as.integer(),
        Nsub2 = df_sum_y2$n %>% as.integer(),
        d1 = df_sum_y1$dose,
        d2 = df_sum_y2$dose,
        Lymean1 =df_logy1$Lmean,
        Lysd1 = df_logy1$LSD,
        ycount2 = df_sum_y2$Incidence
)

# Priors and inits----
priors <- list(
     nu = 3,
     prior_sigma = c(0,0.5),
     power_bound =c(1,18),
     dir1 =c(0,Inf),
     dir2=c(0,1),
     prior_background = c(0,1),
     prior_threshold = c(0,5),
     prior_slope =c(1,1),
     prior_power =c(2,2),
     run_simu = 0
)
#  Compile-----
model_HDH_sum_meta_flat <- cmdstan_model(
     file.path(dir_code,"Hill_DHill_Lognormal_summary_meta_flat.stan")
  )
model_HDH_sum_meta_hier <- cmdstan_model(
     file.path(dir_code,"Hill_DHill_Lognormal_summary_meta_hier.stan")
  )
# Sampling------
model_HDH_sum_meta_flat$sample(
     data = append(datalist_sum,priors),
     iter_warmup = i_warmup,iter_sampling = i_sample,
     output_dir = dir_storage,output_basename = paste0("T",iterID,"
         _fit_HDH_sum_meta_flat")
  )
model_HDH_sum_meta_hier$sample(
     data = append(datalist_sum,priors),
     iter_warmup = i_warmup,iter_sampling = i_sample,
     max_treedepth = 12,
     output_dir = dir_storage,output_basename = paste0("T",iterID,"
         _fit_HDH_sum_meta_hier")
  )
```
